# Elucidating the acid-base mechanisms underlying otolith overgrowth in fish exposed to ocean acidification

**DOI:** 10.1101/2021.11.30.470671

**Authors:** Garfield T. Kwan, Martin Tresguerres

**Affiliations:** Marine Biology Research Division, Scripps Institution of Oceanography, University of California San Diego, USA; NOAA Fisheries Service, Southwest Fisheries Science Center, USA

**Keywords:** Endolymph, climate change, calcification, biomineralization, rockfish, carbon dioxide

## Abstract

Over a decade ago, ocean acidification (OA) exposure was reported to induce otolith overgrowth in teleost fish. This phenomenon was subsequently confirmed in multiple species; however, the underlying physiological causes remain unknown. Here, we report that splitnose rockfish (*Sebastes diploproa)* exposed to ~1,600 μatm *p*CO_2_ (pH ~7.5) were able to fully regulated the pH of both blood and endolymph (the fluid that surrounds the otolith within the inner ear). However, while blood was regulated around pH 7.80, the endolymph was regulated around pH ~8.30. These different pH setpoints result in increased *p*CO_2_ diffusion into the endolymph, which in turn leads to proportional increases in endolymph [HCO_3_^−^] and [CO_3_^2−^]. Endolymph pH regulation despite the increased *p*CO_2_ suggests enhanced H^+^ removal. However, a lack of differences in inner ear bulk and cell-specific Na^+^/K^+^-ATPase and vacuolar type H^+^-ATPase protein abundance localization pointed out to activation of preexisting ATPases, non-bicarbonate pH buffering, or both, as the mechanism for endolymph pH-regulation. These results provide the first direct evidence showcasing the acid-base chemistry of the endolymph of OA-exposed fish favors otolith overgrowth, and suggests that this phenomenon will be more pronounced in species that count with more robust blood and endolymph pH regulatory mechanisms.

## Introduction

The inner ear of teleost fishes contains three pairs of otoliths that contribute to hearing and maintaining balance. Otoliths are comprised of calcium carbonate (CaCO_3_) embedded within a protein matrix, and are biomineralized within an acellular fluid called the endolymph (Payan et al., 2004a). Otoliths are biomineralized in a successive ring pattern correlated with the fish seasonal growth rate [2–4], which are used by scientists and fishery managers to estimate fish age and length [5,6], estimate recruitment, and set fishery-specific catch limits [7,8].

Originally, it was predicted that CO_2_-induced ocean acidification (OA) would impair otolith biomineralization because the associated decreases in seawater pH and [CO_3_^2−^] hamper CaCO_3_ precipitation [9]. However, subsequent studies reported that fish exposed to OA developed enlarged otoliths [10–16]. These findings led to a broader awareness otolith biomineralization is strongly linked to endolymph and blood chemistries, and to the hypothesis that biological regulation of endolymph pH could lead to increased [CO_3_^2−^] resulting in otolith overgrowth [10]. In addition, fish exposed to hypercapnia typically accumulate [HCO_3_^−^] in their plasma to compensate the respiratory acidosis; this could result in enhanced HCO_3_^−^ flux into the endolymph and further contribute to otolith overgrowth [17]. However, experimental support for these hypotheses is lacking, as there are no reports of endolymph acid-base parameters under OA-relevant conditions, and only a few studies have measured blood acid-base parameters in fish exposed to OA-relevant CO_2_ levels [18–20]. This knowledge gap is in large part due to the disrupting effects of regular blood sampling methods on the acid-base status of fish internal fluids, coupled with the difficulty of collecting sufficient endolymph for analyses. Moreover, the cellular heterogeneity of the inner ear complicates the quantification of ionocyte-specific responses using standard molecular and biochemical assays on bulk tissue. As a result, the underlying acid-base and physiological causes of OA-induced otolith overgrowth remain unknown.

The chemistry of the endolymph is actively controlled by the inner ear epithelium to maintain acid-base conditions that promote biomineralization, namely, higher pH, [HCO_3_^−^], [CO_3_^2−^], and total CO_2_ than the blood [21–24]. This gradient is actively maintained by two types of ion-transporting cells (“ionocytes”): the Type-I ionocyte, which transports K^+^ and Cl^−^ into the endolymph and removes H^+^ powered by Na^+^/K^+^-ATPase (NKA) [21,25–27] and the Type-II ionocyte, which secretes HCO_3_^−^ into the endolymph driven by V-type H^+^-ATPase (VHA) [21,25–29]. However numerous other cells within the inner ear organ also express NKA and VHA, including the sensory hair cells and the endothelial cells that make up the blood vessels [26,27,30].

In the current study, splitnose rockfish (*Sebastes diploproa)* were exposed to ~1,600 μatm CO_2_ (pH ~7.5), a condition readily experienced in their natural habitat [31,32] and predicted for the surface ocean by the year 2300 [33]. The OA exposure spanned three days, a duration previously documented to result in otolith overgrowth [16]. Blood acid-base chemistry was measured after taken samples using a benzocaine-based anesthetic protocol that yields measurements comparable to those achieved using cannulation [20]. Additionally, we took advantage of the large rockfish inner ear organ to collect sufficient endolymph for acid-base chemistry analysis, and inner ear tissue for quantification of NKA and VHA protein abundances. Finally, we performed immunohistochemical analyses on six inner ear cell types to explore potential cell-specific changes in protein expression patterns. This multidimensional approach allowed us to explore the mechanistic acid-base causes that underlie otolith overgrowth in fish exposed to OA.

## Methods

### Specimens

Juvenile splitnose rockfish (*S. diploproa*) were caught from drifting kelp paddies off the shores of La Jolla and raised in the Hubbs Experimental Aquarium (La Jolla, USA) in accordance to the permit (#SCP13227) issued by the California Department of Fish and Wildlife. Rockfish were raised for >2 years in a flow-through system with seawater continuously pumped from the Scripps Coastal Reserve, and were fed frozen market squids and food pellets (EWOS, Cargill Incorporated, Minneapolis, MN, USA). Average rockfish total length (11.88 ± 0.29 cm) and weight (42.86 ± 2.79 g) (N=24) were not significantly different between treatments. All experiments were approved under the Institutional Animal Care and Use Committee protocol (#S10320) by the Scripps Institution of Oceanography, University of California San Diego animal care committee.

### Experimental Aquarium Setup

Two header tanks were supplied with ambient seawater from the Scripps Coastal Reserve, one was not manipulated and was considered as the control condition. The other header tank was bubbled with CO_2_ using a pH-stat system (IKS Aquastar, Karlsbad, Germany) to maintain a seawater pH ~7.5 and generate the OA condition. Temperature and pH were continuously monitored and recorded every 2 minutes using the IKS Aquastar system (figure S1). Discrete seawater samples were collected from header tanks at the beginning and end of each experiment, and analysed for alkalinity (via titration with LabView software Version 2.9j; National Instruments, Austin, Texas, United States), pH (using the indicator dye purified m-cresol purple [34] in an Agilent 8453 spectrophotometer (Agilent, Santa Clara, CA, USA)), and salinity (by converting density measurements using Mettler Toledo DE-45 (Mettler-Toledo, Columbus, Ohio, United States)) by the Dickson Lab (Scripps Institution of Oceanography). The pH values from the discrete seawater samples were used to validate and back-correct the IKS pH measurements. Subsequently, the pH, alkalinity, and salinity values were used to calculate *p*CO_2_ using CO2SYS [35]. These analyses indicated control pH and *p*CO_2_ levels of 7.89 ± 0.012 and 571.90 ± 4.88 μatm, respectively, which are typical for La Jolla, USA [36–38]. In contrast, pH and *p*CO_2_ in the OA treatment were 7.49 ± 0.01 and 1,591.56 ± 18.58 μatm, respectively (table S1).

Each header tank supplied water to three opaque 3-L experimental tanks at a flow rate of 0.3-L min^−1^. Individual rockfish were acclimated within an experimental tank for 12 hours, followed by a 72-hour exposure to control or OA conditions. To ensure similar metabolic state among individuals, rockfish were not fed during the 48 hours prior to the acclimations or during the experiment. Three separate experiments were conducted during March 2020, each time with three control and three OA-exposed fish. No mortality was observed.

### Blood, endolymph, and inner ear sampling

Sampling and acid-base determinations were performed in a temperature-controlled room at 18°C (i.e. same as that of seawater). Fish were anesthetized by stopping the seawater flow into the individual experimental tank and slowly adding benzocaine through to achieve a final concentration of 0.15 g/L. After fish lost equilibrium (~5 minutes), they were moved to a surgery table where the gills were irrigated with aeriated seawater containing benzocaine (0.05 g/L) using a pump [20]. Blood was drawn from the caudal vein using a heparinized syringe and pH was immediately measured using a microelectrode (Orion™ PerpHecT™ Ross™, ThermoFisher Scientific, Waltham, MA, USA). Next, blood was centrifuged for 1 minute at 6,000xg using a microcentrifuge (VWR Kinetic Energy 26 Joules, Radnor, PA, USA), and the resulting plasma was measured for total CO_2_ (TCO_2_) using a carbon dioxide analyser (Corning 965 carbon dioxide analyser, Ciba Corning Diagnostic, Halstead, Essex, United Kingdom). After blood sampling (N=8-9), the fish was euthanized by spinal pithing, and the gills were quickly removed. Endolymph (N=7-8) was drawn using a heparinized syringe from the ventral side of the skull, and pH and TCO_2_ were measured as described above. Inner ear tissue was either flash frozen in liquid nitrogen and stored at −80°C, or fixed in 4% paraformaldehyde (8 hours at 4°C), incubated in 50% ethanol (8 hours at 4°C), then stored in 70% ethanol until processing. The sampling procedure, from spinal pithing to endolymph sampling, took less than 3 minutes.

### HCO_3_^−^, CO_3_^2−^, and pCO_2_ calculation

Blood and endolymph pH and TCO_2_ values were used to calculate [HCO_3_^−^], [CO_3_^2−^], and *p*CO_2_ using the Henderson-Hasselbalch equation. The solubility coefficient of CO_2_ (plasma: 0.0578 mmol L^−1^ Torr^−1^; endolymph: 0.0853 mmol L^−1^ Torr^−1^), ionic strength (plasma: 0.15 mol L^−1^, endolymph: 0.18 mol L^−1^), pK_1_^′^ (plasma: ~6.20, endolymph: ~6.16), and pK_2_^′^ (plasma: ~9.76, endolymph: ~9.71) were based upon [39] and [24] for blood and endolymph, respectively. The [Na^+^] (plasma: 170 mmol L^−1^, endolymph: 100 mmol L^−1^) used for calculating pK_1_^′^ was based upon [21].

### Antibodies

NKA was immunodetected using a monoclonal α5 mouse antibody raised against the α-subunit of chicken NKA (a5, Developmental Studies Hybridoma Bank, Iowa City, IA, USA; [40]), whereas the β-subunit of VHA was immunodetected using a custom-made polyclonal rabbit antibody (epitope: AREEVPGRRGFPGYC; GenScript, Piscataway, USA). These antibodies have been previously used in the inner ear of the Pacific chub mackerel (*Scomber japonicus*; [26]), and were validated here for splitnose rockfish (figure S2). Secondary antibodies goat anti-mouse HRP-linked secondary antibodies (Bio-Rad, Hercules, CA, USA) and goat anti-rabbit HRP-linked secondary antibodies (Bio-Rad) were used for immunoblotting.

### Western Blotting and Relative Protein Abundance Analysis

Frozen inner ear samples were immersed in liquid nitrogen, pulverized using a handheld motorized homogenizer (Kimble®/Kontes, Dusseldorf, Germany), and suspended in ice-cold homogenization buffer containing protease inhibitors (250 mmol l^−1^ sucrose, 1 mmol l^−1^ EDTA, 30 mmol l^−1^ Tris, 10 mmol l^−1^ benzamidine hydrochloride hydrate, 1 mmol l^−1^ phenylmethanesulfonyl fluoride, 1 mmol l^−1^ dithiothreitol, pH 7.5). Samples were centrifuged at low speed (3,000xg, 10 minutes, 4°C) to remove debris, and the resulting supernatant was considered the crude homogenate. A subset of this crude homogenate was further centrifuged (21,130xg, 30 minutes, 4°C), and the pellet was saved as the membrane-enriched fraction. Total protein concentration in all fractions was determined by the Bradford assay [41]. Prior to SDS-electrophoresis, samples were mixed with an equal volume of 90% 2x Laemmli buffer and 10% β-mercaptoethanol, and heated at 70°C for 5 minutes. Proteins (crude homogenate: 10 μg per lane; membrane-enriched fraction: 5 μg per lane) were loaded onto a 7.5% polyacrylamide mini gel (Bio-Rad, Hercules, CA, USA) – alternating between control and high CO_2_ treatments to avoid possible gel lane effects. The gel ran at 200 volts for 40 minutes, and the separated proteins were then transferred to a polyvinylidene difluoride (PVDF) membrane using a wet transfer cell (Bio-Rad) at 100 mAmps at 4°C overnight. PVDF membranes were incubated in tris-buffered saline with 1% tween (TBS-T) with milk powder (0.1 g/mL) at RT for 1 hour, then incubated with primary antibody (NKA: 10.5 ng/ml; VHA: 3 μg/ml) in blocking buffer at 4°C overnight. On the following day, PVDF membranes were washed with TBS-T (three times; 10 minutes each), incubated in blocking buffer with anti-rabbit secondary antibodies (1:10,000) at RT for 1 hour, and washed again with TBS-T (three times; 10 minutes each). Bands were made visible through addition of ECL Prime Western Blotting Detection Reagent (GE Healthcare, Waukesha, WI) and imaged and analysed in a BioRad Universal III Hood using Image Lab software (version 6.0.1; BioRad). Following imaging, the PVDF membrane was incubated in Ponceau stain (10 minutes, room temperature) to estimate protein loading. Relative protein abundance (N=6-8) were quantified using the Image Lab software (version 6.0.1; BioRad) and normalized by the protein content in each lane.

### Whole-mount immunohistochemistry and confocal microscopy

Immunolabeling was performed based on the protocol described in Kwan *et al.*, (2020) for tissue sections and optimized for whole tissues as follows. Fixed inner ear tissue was rehydrated in phosphate buffer saline + 0.1% tween (PBS-T) for 10 min. Autofluorescence was quenched by rinsing in ice-cold PBS-T with sodium borohydride (1.5 mg/mL; six times; 10 minutes each), followed by incubation in blocking buffer (PBS-T, 0.02% normal goat serum, 0.0002% keyhole limpet hemocyanin) at room temperature for one hour. Samples were incubated with blocking buffer containing primary antibodies (NKA: 40 ng/mL; VHA: 6 μg/mL) at 4°C overnight. On the following day, samples were washed in PBS-T (three times at room temperature; 10 minutes each), and incubated with the fluorescent secondary antibodies (1:500) counterstained with DAPI (1 μg/mL) at room temperature for 1 hour. Samples were washed again in PBS-T as before and stored at 4°C until imaging.

Immunostained inner ear samples were immersed in PBS-T, mounted onto a depressed glass slide fitted with a glass cover slip (No. 1.5, 0.17 mm) and imaged using a Zeiss LSM800 inverted confocal microscope equipped with a Zeiss LD LCI Plan-Apochromat 40x/1.2 lmm Korr DIC M27 objective and Zeiss ZEN 2.6 blue edition software (Cambridge, United Kingdom). The following channels were used for imaging: VHA (excitation 493 nm with 1% laser power, emission 517 nm, detection 510– 575 nm), NKA (excitation 577 nm at 1% laser power, emission 603 nm, detection 571–617 nm), and DAPI (excitation 353 nm at 0.7% laser power, emission 465 nm, detection 410–470 nm). Z-stacks (range: ~70–400 optical sections; thickness: ~0.27 μm per section) of the various inner ear cell types were visualized as maximum intensity projection, and through orthogonal cuts to capture fluorescent signal across the X-Z and Y-Z planes. Inner ear organs from four control and four OA-exposed rockfish were imaged.

### Statistical Analysis

Normality was tested using the Shapiro-Wilk normality test, and homogeneity was tested using the F-test. Datasets that failed to meet the assumptions of normality were log-(i.e. [CO_3_^2−^], pH) or inverse-transformed (i.e. [H^+^]). Acid-base parameters were analysed using two-way analysis of variance (2-way ANOVA), with “CO_2_ level” (control or OA) and “internal fluid” (blood or endolymph) as factors. If significant interaction effect was detected, subsequent Tukey honest significant difference (HSD) tests were used. NKA and VHA protein abundances were analysed using two-tailed Student’s t-tests. Values are reported as mean ± s.e.m., and an alpha of 0.05 was employed for all analyses. Statistical tests were performed using Prism (version 7.0a) and R (version 4.0.3; R Development Core Team, 2013).

## Results and discussion

The difference in seawater *p*CO_2_ between the control and OA-condition was ~1,000 μatm, which induced an equivalent elevation in blood *p*CO_2_ from 1,603.25 ±190.69 μatm in control fish to 2,659.20 ± 223.87 μatm in OA-exposed fish (figure 1a). However, blood pH was fully regulated (control: 7.75 ± 0.03; OA: 7.85 ± 0.04) (figure 1b). As is typical for regulation of blood acidosis [43], OA-exposed fish demonstrated a significant accumulation of HCO_3_^−^ in blood plasma, from 2.37 ± 0.20 mM in control fish up to 5.16 ± 0.31 mM in OA-exposed fish (figure 1c). This response matches the magnitude of the hypercapnic stress according to classic Davenport acid-base physiology, as well as the three previous studies on blood acid-base chemistry in fish exposed to OA-relevant CO_2_ levels [18–20]. In addition, the increased plasma TCO_2_ at unchanged pH led to the tripling of plasma [CO_3_^2−^] from ~0.02 to ~0.07 mM (figure 1d). These increases in blood [HCO_3_^−^] and [CO_3_^2−^] may contribute to the skeletal hypercalcification [44] and deformities [45] reported in some OA-exposed fishes.

**Figure 1:**
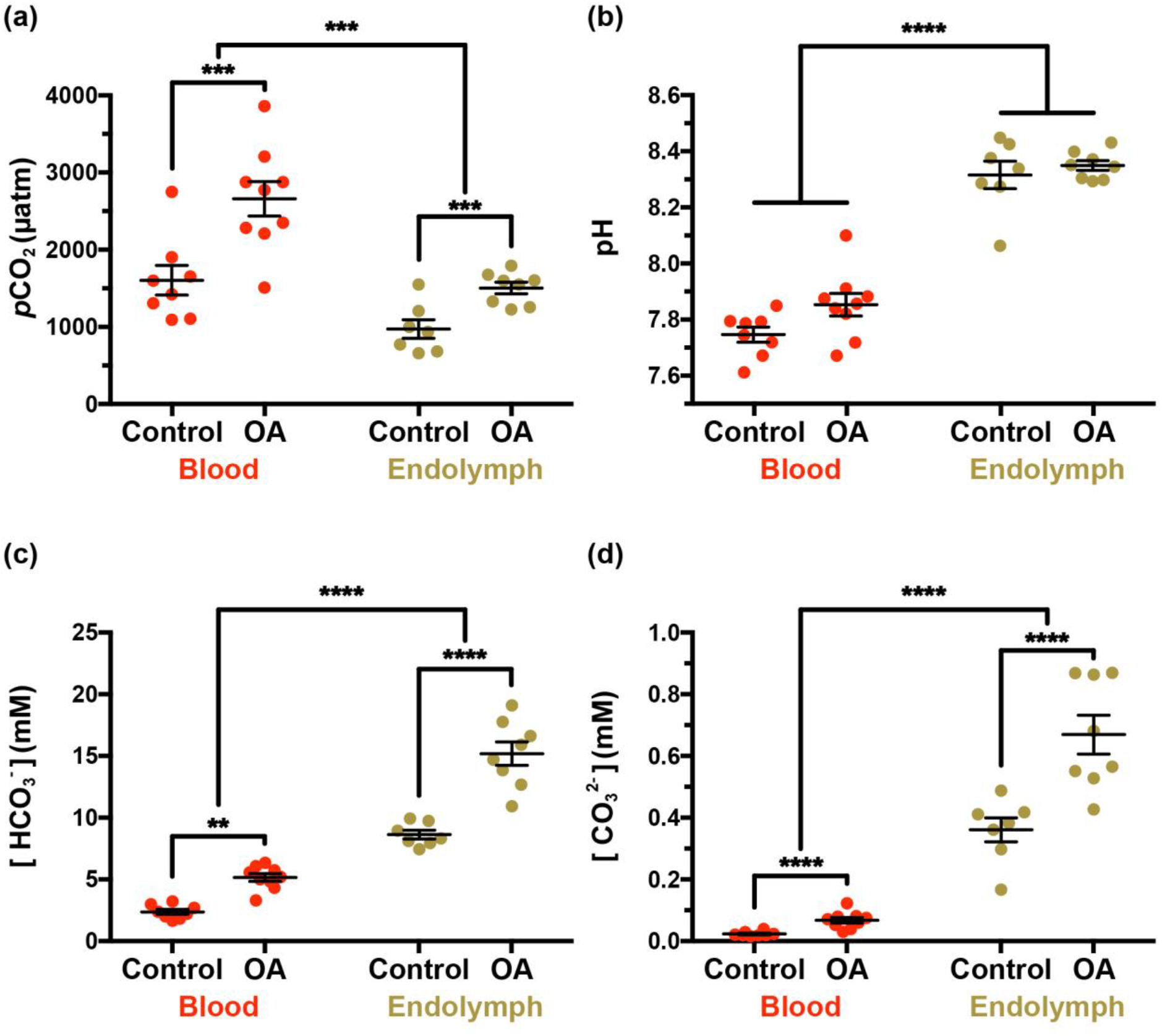
Blood and endolymph acid-base parameters in control and OA-exposed rockfish. **A)** *p*CO_2_, **B)** pH, **C)** [HCO_3_^−^], and **D)** [CO_3_^2−^]. Data is presented as mean and s.e.m. for each group and the individual measurements are shown as red (blood) or beige (endolymph) points (N= 7-9). Statistical significance between fluids, and between treatments for a given fluid are indicated by the connecting lines and asterisks (2-way ANOVA, *p<0.05, **p<0.005, ***p<0.001, ****p<0.0001). Statistical details are reported in tables S3-S5, and TCO_2_,[CO_2_] and [H^+^] are shown in figure S3.

The endolymph of control rockfish had higher TCO_2_ compared to the blood (9.06 ± 0.38 vs 2.46 ± 0.20 mM; figure S3) and also a higher pH (8.32 ± 0.05 vs. 7.75 ± 0.03), resulting in lower *p*CO_2_ (971.38 ± 120.70 vs. 1,603.25 ± 190.69 μatm), higher [HCO_3_^−^] (8.63 ± 0.35 vs. 2.37 ± 0.20 mM), and much higher [CO_3_^2−^] (0.36 ± 0.04 vs 0.02 ± 0.01 mM) (figure 1a-d). Importantly, these measurements revealed higher pH and lower pCO_2_, TCO_2_, [HCO_3_^−^] and [CO_3_^2−^] compared to previous studies that collected endolymph without previously anesthetizing the fish [21,22], and to others that used 2-phenoxyethanol as anesthetic but did not irrigate the gills during endolymph collection [23,24] (table S2). This finding highlights the crucial importance of sampling procedures for accurate acid-base measurements in fish physiological fluids. Indeed, fish struggling during handling and hypoxia due to gill collapse during emersion are known to greatly affect blood acid-base measurements, and our results indicate that these disturbances extend to the endolymph.

In rockfish exposed to OA, endolymph *p*CO_2_ increased from 971.38 ± 120.70 to 1503.21 ± 73.72 μatm (figure 1a). Crucially, this ~500 μatm increase was half of that observed in the blood and therefore the *p*CO_2_ difference between blood and endolymph increased from ~600 to ~1,100 μatm, which is predicted to induce a proportional increase in CO_2_ flux into the endolymph following Fick’s law of diffusion. Endolymph TCO_2_ in OA-exposed rockfish also nearly doubled (control: 9.06 ± 0.38 mM; OA= 15.96 ± 1.02 mM; figure S3) and, since pH remained unchanged at ~8.30 pH (figure 1b), it was reflected as increased [HCO_3_^−^] (control: 8.63 ± 0.35 mM, OA: 15.19 ± 0.95 mM vs) (figure 1c) and [CO_3_^2−^] (control: 0.36 ± 0.04 mM; OA 0.67 ± 0.06 mM) (figure 1d). Since aragonite saturation state (Ω_aragonite_) is directly proportional to [CO_3_^2−^], it implies that biomineralization in the endolymph of OA-exposed fish is nearly twice more favorable than in that of control fish. To our knowledge, this is the first direct evidence that the acid-base chemistry in the endolymph of OA-exposed fish favors otolith overgrowth.

The increased *p*CO_2_ diffusive rate into the endolymph and subsequent generation of H^+^ as a result of CO_2_ hydration and CaCO_3_ biomineralization are bound to induce a decrease in pH. Thus, the lack of change in endolymph pH in OA-exposed rockfish indicates robust pH regulation. Thus, we hypothesized that OA-exposed fish may have increased abundance of NKA and VHA, as these ATPases are proposed to provide the driving force for transepithelial H^+^ and HCO_3_^−^ transport across the inner ear epithelium [21,26,27,30]. However, Western blotting on bulk inner ear tissue revealed no significant differences between control and OA-exposed fish (figure 2, table S6).

**Figure 2:**
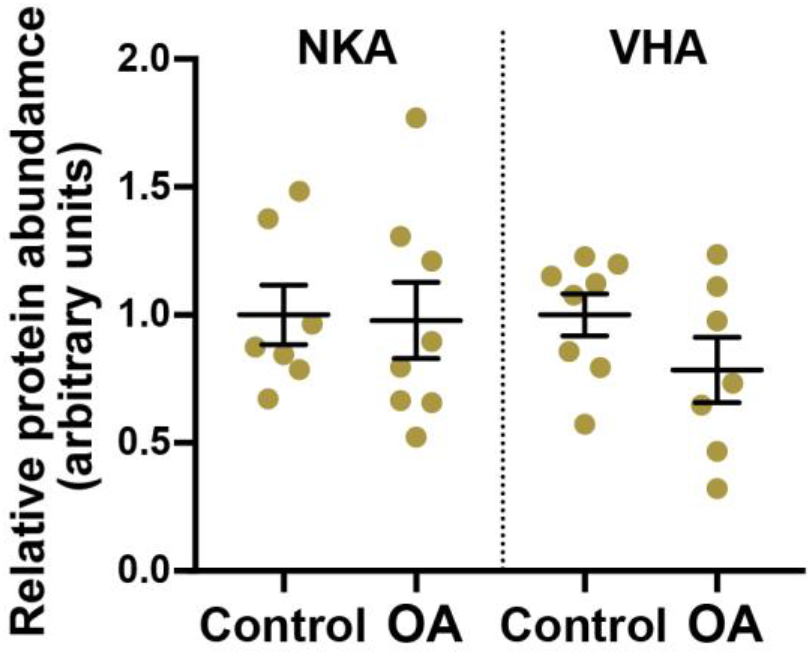
Na^+^/K^+^-ATPase (NKA) and V-type H^+^-ATPase (VHA) protein abundance in the inner ear organ of control and OA-exposed rockfish. Data is presented as mean and s.e.m. and the individual measurements are shown as beige points (N= 7-8). Relative protein abundance was calculated for each ATPase; NKA and VHA abundances are not comparable to each other. There were no significant differences for NKA (*p*=0.9104) or VHA (*p*=0.1695). Statistical details are reported in tables S6.

Next, we used immunocytochemistry and confocal microscopy to examine potential changes in NKA and VHA abundance or sub-cellular localization in specific inner ear epithelial cell types. The NKA and VHA immunostaining in rockfish inner ear epithelial cells generally matched reports from other fish species [25,26,30,46] (figure 3; figure S4), and there were no apparent differences between control and OA-exposed fish in any cell type in terms of signal intensity of subcellular localization. The Type-I ionocytes are characterized by intense NKA signal in their highly infolded basolateral membrane and by a much fainter cytoplasmic VHA signal (figure 3a). These ionocytes are most abundant in the meshwork area, where they contact each other by their pseudopods giving the appearance of an interconnected matrix. The Type-II ionocytes are interspersed between the Type-I ionocytes in the meshwork area and have cytoplasmic VHA signal of comparable intensity to that in the Type-I ionocytes; however, they lack NKA signal (figure 3a). The sensory hair cells are in the macula; they express intense NKA signal in their basolateral membrane and very intense cytoplasmic VHA signal, which was especially concentrated towards their basal area consistent with synaptic vesicles (figure 3b). The supporting cells surround each sensory hair cell; they display faint cytoplasmic VHA signal and no detectable NKA signal (figure 3b). The granular cells flank the macula and have a characteristic columnar shape. These cells have faint NKA signal along their lateral plasma membrane and faint cytoplasmic VHA signal (figure 3c). Finally, the squamous cells are found in the patches area in the distal side of the epithelium; these cells are very thin and have NKA signal on their ribbon-like lateral membrane as well as faint cytoplasmic VHA signal (figure 3d). A summary of the NKA and VHA relative signal intensities in each cell type is reported in table S7.

**Figure 3:**
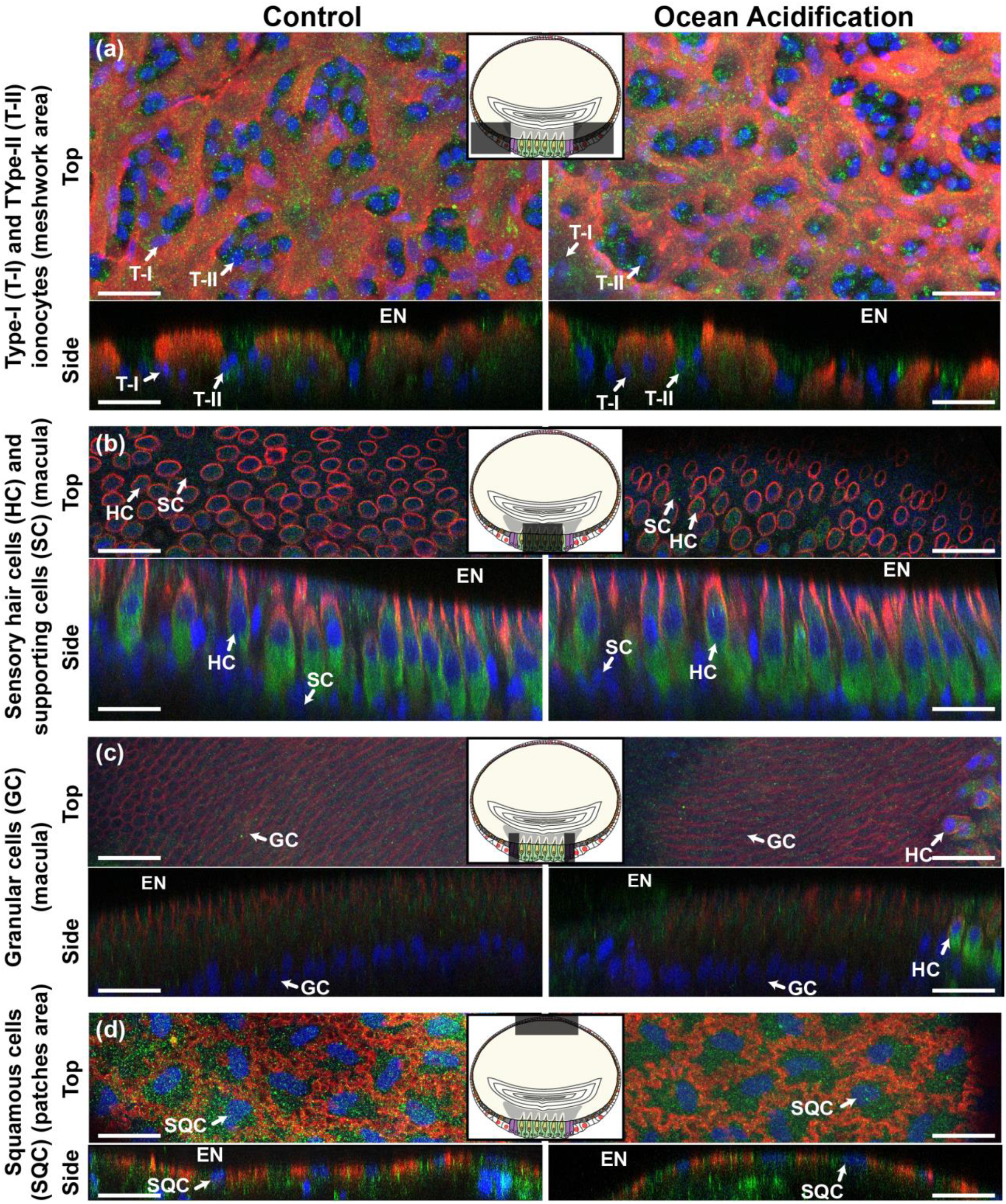
Immunocytochemistry of the inner ear epithelium of control and OA-exposed rockfish. Na^+^/K^+^-ATPase is in red, V-type H^+^-ATPase is in green, and nuclei are in blue. There were no apparent differences in NKA or VHA signal intensities or localization patterns between control and OA-exposed fish. **(a)** Type-I (T-I) and Type-II ionocytes (T-II), **(b)** sensory hair cells (HC) and supporting cells (SC), **(c)** granular cells (GC), and **(d)** squamous cells (SQC). The top view shows the X-Y plane in maximum projection, whereas the side view shows the X-Z or Y-Z plane using orthogonal cuts. EN = endolymph. Scale bar = 20 μm. Images are representative of inner ear from four control and four OA-exposed rockfish. The shaded boxes in the diagrams indicate the location of each cell type within the otolith sac. A larger diagram showcasing the heterogeneous cellular anatomy of the inner ear epithelium is provided in figure S4.

The lack of apparent differences in NKA and VHA abundance and localization cellular patterns between control and OA-exposed fish indicates that preexisting levels of NKA and VHA were sufficient to mediate the endolymph pH regulation observed in our study. Overall, these findings are consistent with models suggesting that H^+^ extrusion from the endolymph into the blood passively follows the transepithelial potential that is established by active K^+^ excretion into the endolymph [1]. And since the function of the sensory hair cells requires a high [K^+^] in the endolymph, modulation of inner ear transepithelial potential for the sole purpose of decreasing H^+^ extrusion seems unlikely.

In our recent paper [26], we proposed that HCO_3_^−^ transport into the endolymph and H^+^ removal could be upregulated by insertion of VHA into the basolateral membrane of Type-II ionocytes; however, we found no evidence for such mechanism in OA-exposed rockfish (figure 3a, *right panels*). Instead, upregulation of ATPase activity could have occurred *via* other post-translational modifications or by increased substrate availability (c.f. [47]). The expression of carbonic anhydrases, ion exchangers, and other acid-base relevant proteins must be examined in future studies, ideally through an approach that includes cell-specific analyses. Lastly, a contribution of non-bicarbonate buffering to endolymph pH regulation cannot be ruled out; unfortunately, performing the required titrations are not trivial due to the small volume of this fluid.

## Conclusions

Increased endolymph [HCO_3_^−^] and [CO_3_^2−^] provides a mechanistic explanation for otolith overgrowth in OA-exposed fish, a phenomenon that was first described over a decade ago [10]. The ultimate cause is an interplay between blood and endolymph acid-base regulation which results in increased CO_2_ flux into the endolymph coupled with endolymph pH regulation. As a result, the carbonate equilibria reactions shift to the right, promoting [HCO_3_^−^] and [CO_3_^2−^] accumulation bound to increase Ωaragonite, and thus promote biomineralization (figure 4). This implies that otolith overgrowth in response to OA will be more pronounced in fish species with more robust acid-base regulatory mechanisms; however, this hypothesis must be experimentally tested. Future studies should also investigate whether the fish inner ear epithelium can curve otolith overgrowth during prolonged OA exposure; for example, by changing the endolymph pH setpoint, modulating glycoprotein or Ca^2+^ secretion, or engaging other compensatory mechanisms. Coupled with functional studies (e.g. [11,48]), this information will help predict whether the inner ear vestibular and auditory sensory systems of fish will be affected by OA. Furthermore, understanding the mechanisms responsible for otolith biomineralization and overgrowth during OA exposure can help improve the accuracy of otolith-reliant aging techniques in the future ocean.

**Figure 4:**
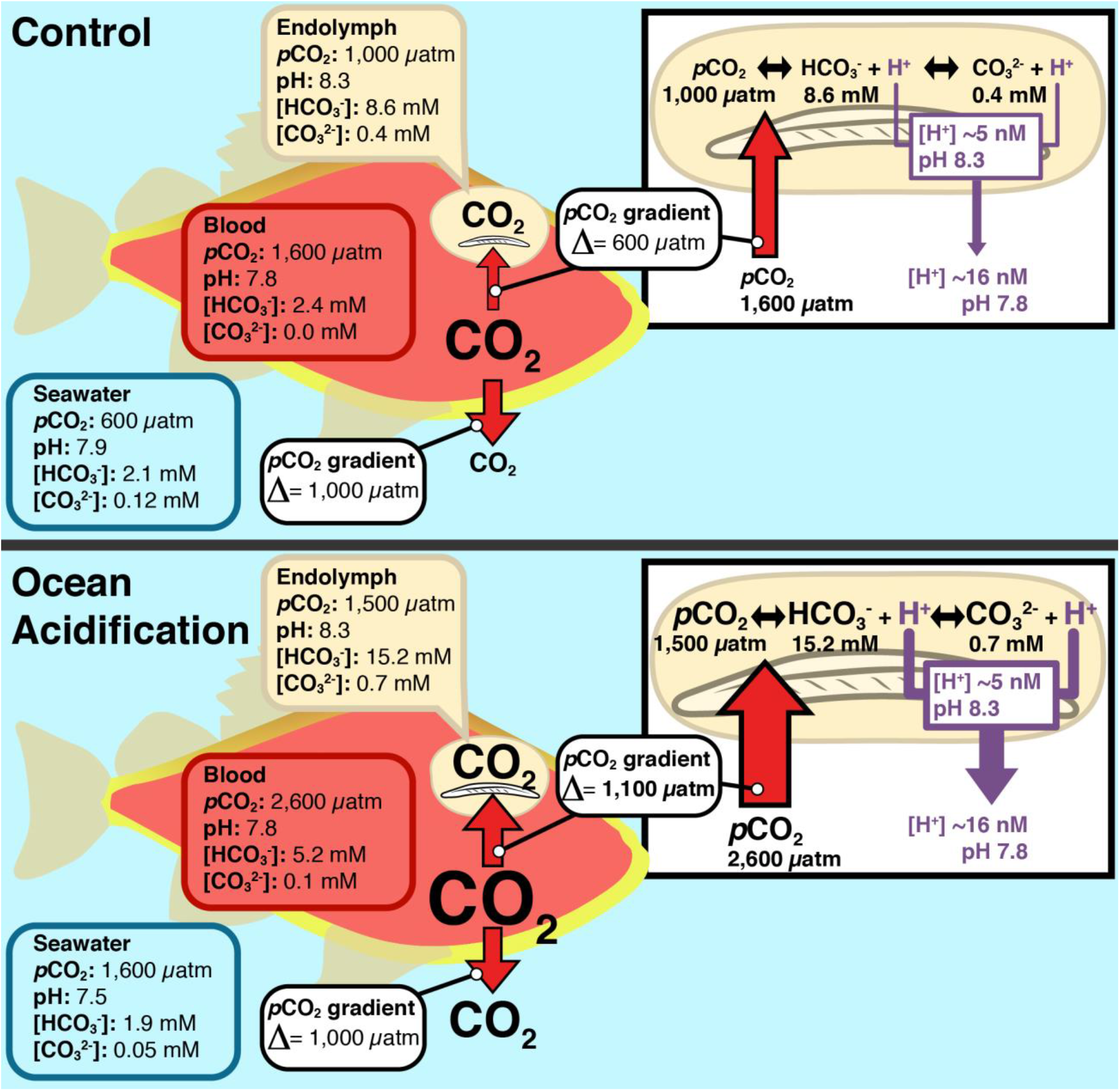
Effect of blood and endolymph acid-base regulation on otolith overgrowth during exposure to ocean acidification. Under control conditions, metabolically produced CO_2_ results in higher levels within the fish blood (~1,600 μatm) than those in seawater (~600 μatm) and endolymph (~1,000 μatm). As a result, blood CO_2_ diffuses into seawater (∆ = ~1,000 μatm) as it passes through the gills, and into the endolymph (∆ = ~600 μatm) as it passes through the inner ear. Under ocean acidification, the 1,000 μatm increase in seawater *p*CO_2_ (to ~1,600 μatm) induces an equivalent increase in the blood (to ~2,600 μatm), but a lesser increase in the endolymph (to ~1,500 μatm). Thus, the *p*CO_2_ diffusion gradient from the blood into seawater remain constant, but the *p*CO_2_ diffusion gradient from the blood into the endolymph increases (∆ = ~1,100 μatm). This process is driven by pH regulation from the endolymph by the inner ear epithelium, presumably by increased H^+^ removal into the blood (although non-bicarbonate buffering cannot be ruled out). The increased CO_2_ diffusion rate into the endolymph coupled with endolymph pH regulation results in the accumulation of [HCO_3_^−^] and [CO_3_^2−^], thereby increasing Ω_aragonite_ and promoting otolith calcification. The size of the arrows is proportional to the fluxes of CO_2_ or H^+^.

## Acknowledgements

This research was supported by National Science Foundation (NSF) grant IOS #1754994 to M.T, and NSF Graduate Research Fellowship and NSF Postdoctoral Research Fellowship in Biology (award #1907334) to GTK. We thank Phil Zerofski (SIO) for collecting the fish and helping with aquarium matters. We are grateful to Taylor Smith, Shane Finnerty, Gabriel Lopez, Kaelan Prime, and Dr. Till Harter for their help with fish care. We also thank Kaelan Prime and Dr. Till Harter for their assistance with experimental maintenance and sampling.

## Author contributions

GTK and MT conceived and designed the experiments, analysed the data, and wrote the manuscript. GTK executed the experiments.

