## Supplemental Materials for "Elucidating the acid-base mechanisms underlying otolith overgrowth in fish exposed to ocean acidification"

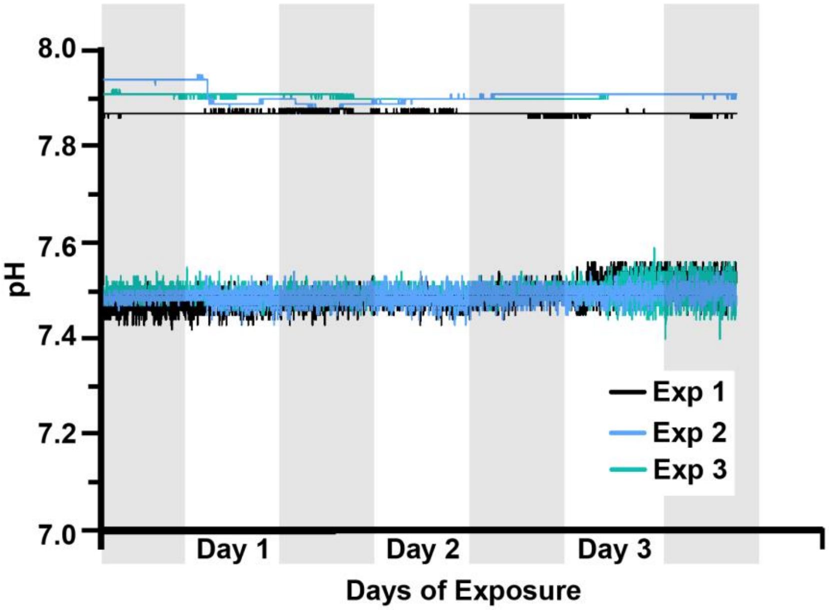


**Figure S1:** Seawater pH in the experimental header tanks, which was monitored and recorded every 2 minutes using an IKS Aquastar system (see Methods for more details).

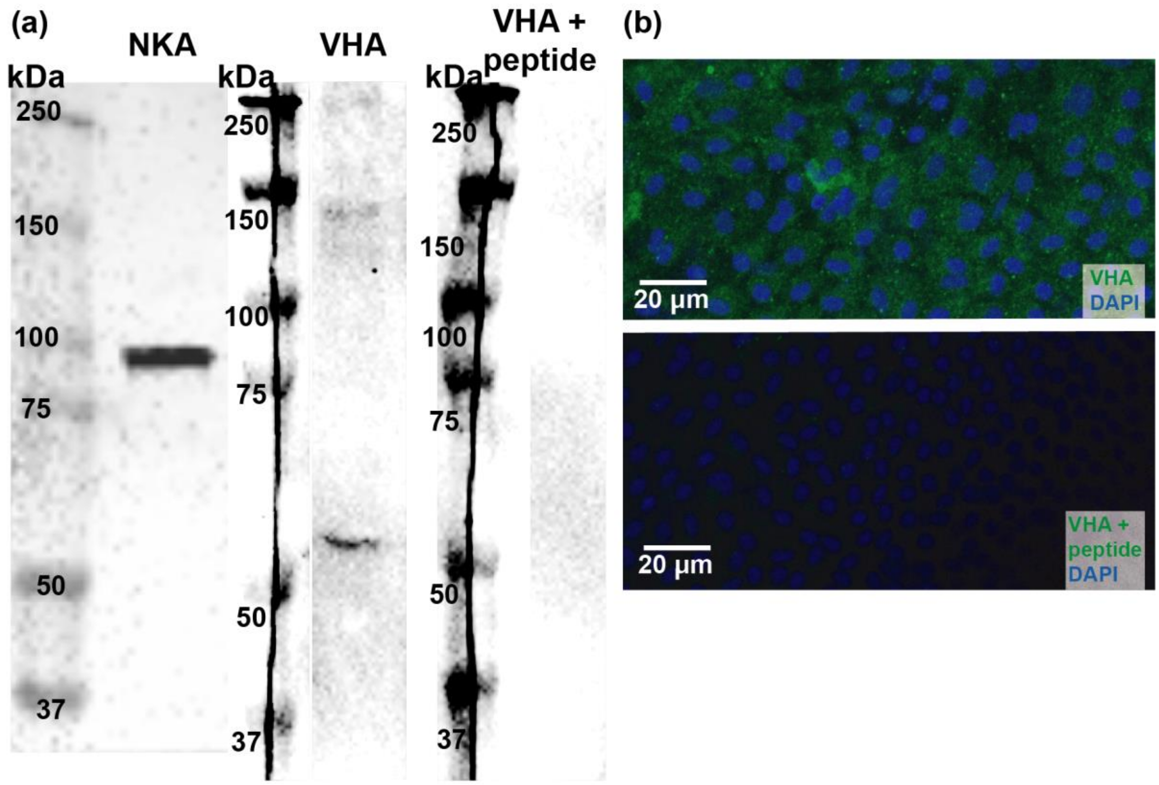


**Figure S2:** Validation of anti-$Na^+$/$K^+$-ATPase (NKA) and vacuolar-type $H^+$-ATPase (VHA) antibodies in rockfish inner ear. **(a)** Western blotting revealed single bands matching the predicted sizes of NKA at ~100 kDa, and VHA at ~55 kDa. The far-right lane shows a lack of VHA signal in the peptide preabsorption control (VHA + peptide), indicating antibody specificity. **(b)** Immunostaining of inner ear tissue showing VHA signal (green) and lack of signal in the peptide preabsorption control. Nuclei were stained with DAPI (blue). In both the Western blot and immunohistochemistry experiments, the anti-VHA antibodies were preincubated with excess antigen peptide (1:5 on a molar basis, overnight, 4°C) and then applied *in lieu* of the primary antibodies (see Methods for more details).

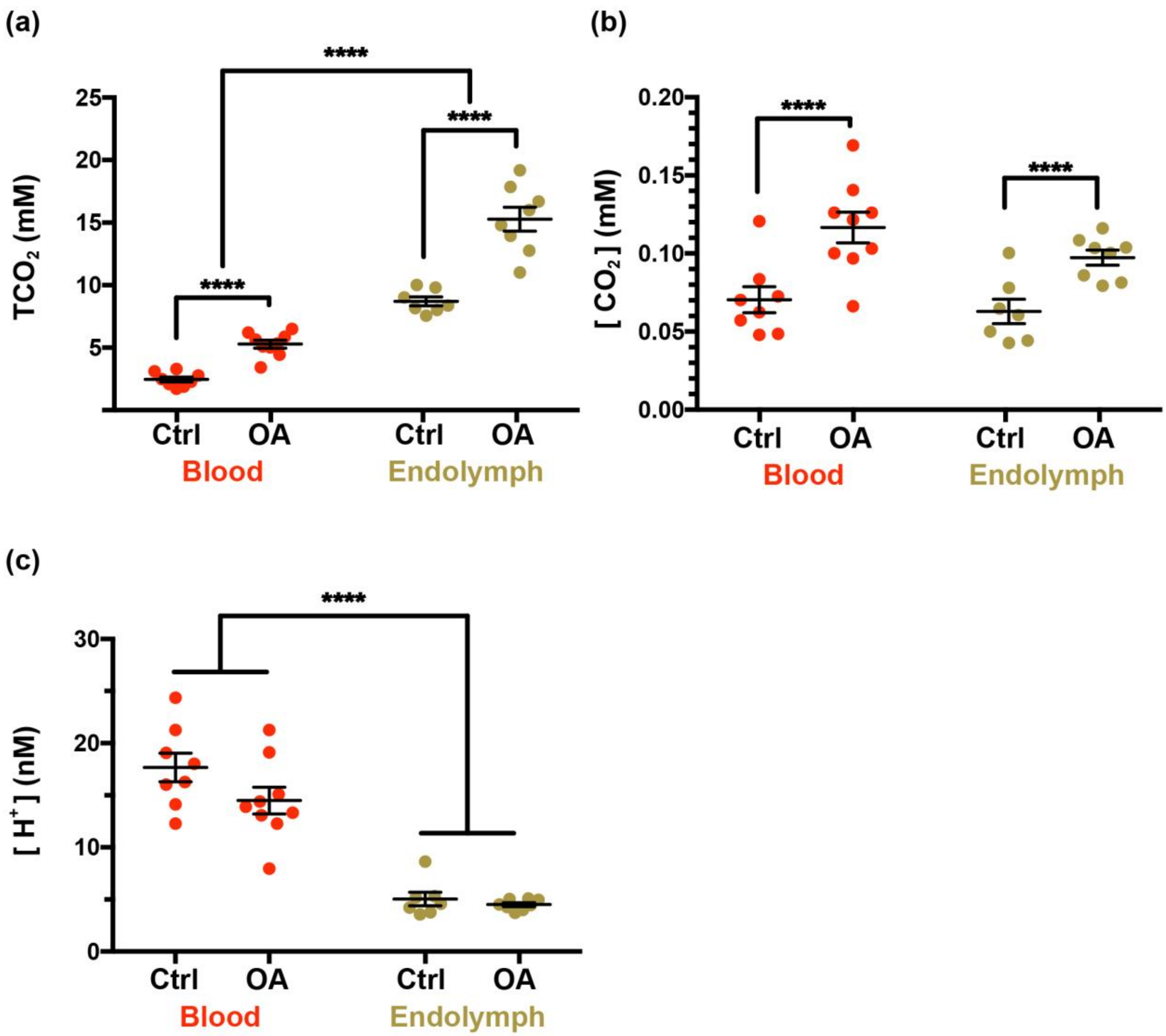


**Figure S3:** Blood and endolymph acid-base parameters in control and OA-exposed rockfish. **a)** [$TCO_2$], **b)** [$CO_2$], **c)** [$H^+$]. Data is presented as mean and s.e.m. for each group and the individual measurements are shown as red (blood) or beige (endolymph) points (N= 7-9). Statistical significance between fluids, and between treatments for a given fluid are indicated by the connecting lines and asterisks (2-way ANOVA, *p<0.05, **p<0.005, ***p<0.001, ****p<0.0001). Statistical details are reported in tables S3-S5.

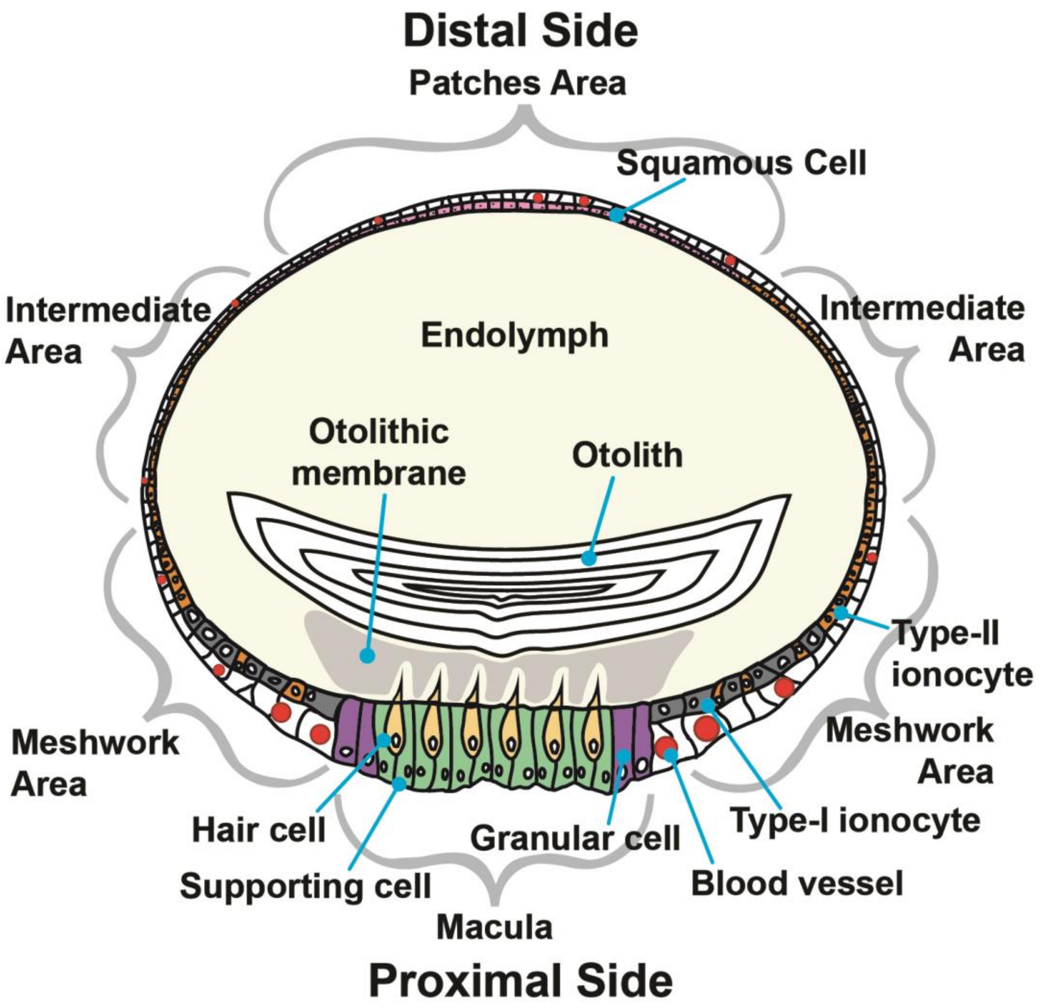


**Figure S4:** Cellular anatomy of the otolith sac inner ear epithelium. The otolith is biomineralized within the endolymph (beige). The sensory hair cells (yellow), supporting cells (green), and granular cells (purple) are found in the macula. The Type-I (dark grey) and Type-II ionocytes (orange) are predominantly present in the meshwork area. Type-II ionocytes make up the intermediate area, and the squamous cells (pink) are found in the patches area. Blood vessels (red) surround the organ, and are especially prevalent around the macula and meshwork areas.

**Table S1**: Seawater parameters of the control and ocean acidification treatments. *Asterisk* and bold font denote a significant difference (Student's T-test, $\alpha$=0.05). Values are mean ± S.E.M calculated from pH recordings at 2-min intervals throughout the exposures (see Methods for details).

| | **Control** | **Ocean Acidification** |
|---|---|---|
| pH | 7.89 ± 0.01 | **7.49 ± 0.01*** |
| Alkalinity (μmol kgSW−1) | 2,229.65 ± 1.62 | 2,230.33 ± 3.27 |
| $pCO_2$ (μatm) | 571.90 ± 4.88 | **1,591.56 ± 18.58 *** |
| Salinity (ppt) | 33.27 ± 0.04 | 33.27 ± 0.04 |
| Temperature (°C) | 15.830 ± 0.003 | |

**Table S2:** Comparison of endolymph acid-base values across studies. *****Values were calculated using data provided in their study. **‡**Values were estimated from charts provided in the study using *WebPlotDigitizer* (version 4.5; apps.automeris.io).

| **Study** | **Payan *et al.* 1997** | | **Takagi 2002** | | **Takagi *et al.* 2005** | | **Kwan & Tresguerres (this study)** | |
|---|---|---|---|---|---|---|---|---|
| **Species** | Rainbow trout (*O. mykiss*) | Turbot (*P. maximus*) | Rainbow trout (*O. mykiss*) | Rainbow trout (*O. mykiss*) | Rainbow trout (*O. mykiss*) | Rainbow trout (*O. mykiss*) | Splitnose rockfish (*S. diploproa*) | Splitnose rockfish (*S. diploproa*) |
| **Anesthesia** | *None reported* | | 0.1% 2-phenoxy-ethanol | | 0.1% 2-phenoxy-ethanol | | 0.05 g/L benzocaine | |
| **Gill Irrigation** | no | | no | | no | | yes | |
| **Treatment** | None | None | 1+ year old | 2+ year old | Day | Night | Control | OA-exposed |
| **pH** | 7.97 ± 0.045 | *n/a* | 7.90 ± 0.02 | 7.95 ± 0.03 | 7.73 ± 0.05**‡** | 7.81 ± 0.04**‡** | 8.32 ± 0.05 | 8.35 ± 0.02 |
| **$TCO_2$ (mM)** | 32.3 ± 2.60 | 17.2 ± 1.04 | *n/a* | *n/a* | *n/a* | *n/a* | 9.06 ± 0.38 | 15.96 ± 1.02 |
| **$pCO_2$ (µatm)** | 8,319 ± 525***** | *n/a* | 16,421 ± 1,013 | 11,078 ± 316 | 15,168 ± 850**‡** | 12,838 ± 980**‡** | 971.38 ± 120.70 | 1,503.21 ± 73.72 |
| **$[HCO_3^-]$ (mM)** | 31.76 ± 5.23***** | *n/a* | 44.6 ± 1.8 | 35.4 ± 2.3 | 27.5 ± 3.5**‡** | 28.2 ± 3.1**‡** | 8.63 ± 0.35 | 15.19 ± 0.95 |
| **$[CO_3^{2-}]$ (mM)** | 0.58 ± 0.23***** | *n/a* | 0.691 ± 0.039 | 0.673 ± 0.077 | 0.30 ± 0.07**‡** | 0.36 ± 0.07**‡** | 0.36 ± 0.04 | 0.67 ± 0.06 |

**Table S3:** Splitnose rockfish blood plasma and endolymph acid-base values following a 3-day exposure to control or ocean acidification (OA) conditions. Control = ~600 µatm $pCO_2$, pH ~7.9; OA = ~1,600 µatm $pCO_2$, pH ~7.5. Values are mean ± s.e.m.

| | **Blood Plasma** | | **Endolymph** | |
|---|---|---|---|---|
| | **Control** | **OA** | **Control** | **OA** |
| **$pCO_2$ (µatm)** | 1,603.25 ± 190.69 | 2,659.20 ± 223.87 | 971.38 ± 120.70 | 1503.21 ± 73.72 |
| **$[CO_2]$ (mM)** | 0.07 ± 0.01 | 0.12 ± 0.01 | 0.06 ± 0.01 | 0.10 ± 0.00 |
| **$[H^+]$ (nM)** | 17.69 ± 1.38 | 14.50 ± 1.29 | 5.04 ± 0.65 | 4.50 ± 0.18 |
| **pH** | 7.75 ± 0.03 | 7.85 ± 0.04 | 8.32 ± 0.05 | 8.35 ± 0.02 |
| **$[HCO_3^-]$ (mM)** | 2.37 ± 0.20 | 5.16 ± 0.31 | 8.63 ± 0.35 | 15.19 ± 0.95 |
| **$[CO_3^{2-}]$ (mM)** | 0.02 ± 0.00 | 0.07 ± 0.01 | 0.36 ± 0.04 | 0.67 ± 0.06 |
| **$TCO_2$ (mM)** | 2.46 ± 0.20 | 5.34 ± 0.32 | 9.06 ± 0.38 | 15.96 ± 1.02 |
| **n** | 8 | 9 | 7 | 8 |

**Table S4:** Two-way analysis of variance (ANOVA; $\alpha$=0.05) on the acid-base chemistry of splitnose rockfish internal fluids (blood, endolymph) after a 3-day exposure to control (~600 µatm $pCO_2$) or ocean acidification (~1,600 µatm $pCO_2$) condition. Bolded values denote significant difference.

| | | Df | Sum Sq | Mean Sq | F-ratio | *p*-value |
|---|---|---|---|---|---|---|
| $pCO_2$ | Fluid | 1 | 6342574 | 6342574 | 27.07 | **<0.0001** |
| | Treatment | 1 | 5002441 | 5002441 | 21.35 | **<0.0001** |
| | Fluid*Treatment | 1 | 545068 | 545068 | 2.326 | 0.1384 |
| | Residual | 28 | 6561019 | 234322 | | |
| $[CO_2]$ | Fluid | 1 | 0.001429 | 0.001429 | 2.715 | 0.1106 |
| | Treatment | 1 | 0.01297 | 0.01297 | 24.64 | **<0.0001** |
| | Fluid*Treatment | 1 | 0.000282 | 0.0002815 | 0.53 | 0.4707 |
| | Residual | 28 | 0.01474 | 0.0005264 | | |
| $[H^+]$ | Fluid | 1 | 0.1857 | 0.1857 | 182.1 | **<0.0001** |
| | Treatment | 1 | 0.001289 | 0.001289 | 1.264 | 0.2705 |
| | Fluid*Treatment | 1 | 0.000038 | 0.000038 | 0.0373 | 0.8482 |
| | Residual | 28 | 0.02855 | 0.00102 | | |
| **pH** | Fluid | 1 | 0.2289 | 0.2289 | 228.3 | **<0.0001** |
| | Treatment | 1 | 0.003157 | 0.003157 | 3.15 | 0.0868 |
| | Fluid*Treatment | 1 | 0.0006252 | 0.0006252 | 0.6237 | 0.4363 |
| | Residual | 28 | 0.02807 | 0.001002 | | |
| $[HCO_3^-]$ | Fluid | 1 | 526.8 | 526.8 | 225.5 | **<0.0001** |
| | Treatment | 1 | 173.1 | 173.1 | 74.09 | **<0.0001** |
| | Fluid*Treatment | 1 | 28.16 | 28.16 | 12.06 | **0.0017** |
| | Residual | 28 | 65.41 | 2.336 | | |
| $[CO_3^{2-}]$ | Fluid | 1 | 50.66 | 50.66 | 434.1 | **<0.0001** |
| | Treatment | 1 | 5.484 | 5.484 | 47 | **<0.0001** |
| | Fluid*Treatment | 1 | 0.3148 | 0.3148 | 2.698 | 0.1117 |
| | Residual | 28 | 3.267 | 0.1167 | | |
| $TCO_2$ | Fluid | 1 | 525.1 | 525.1 | 222.9 | **<0.0001** |
| | Treatment | 1 | 176.1 | 176.1 | 74.75 | **<0.0001** |
| | Fluid*Treatment | 1 | 27.99 | 27.99 | 11.88 | **0.0018** |
| | Residual | 28 | 65.97 | 2.356 | | |

**Table S5:** Tukey HSD posthoc analysis (95% family-wise confidence level; $\alpha$=0.05) following the two-way ANOVA analysis on the acid-base chemistry of splitnose rockfish internal fluids (blood, endolymph) after a 3-day exposure to control (ctrl; ~600 µatm $pCO_2$) or ocean acidification (OA; ~1,600 µatm $pCO_2$) condition. Bolded values denote significant differences.

| | Treatment:Fluid | Difference | Lower Interval | Upper Interval | ***Adjusted p*-value** |
|---|---|---|---|---|---|
| **$[HCO_3^-]$** | Ctrl:Endo – Ctrl:Blood | 6.263 | 4.104 | 8.423 | **<0.0001** |
| | OA:Blood – Ctrl:Blood | 2.786 | 0.7585 | 4.814 | **0.0043** |
| | OA:Endo – Ctrl:Blood | 12.82 | 10.73 | 14.9 | **<0.0001** |
| | OA:Blood – Ctrl:Endo | -3.477 | -5.58 | -1.374 | **0.0006** |
| | OA:Endo – Ctrl:Endo | 6.554 | 4.394 | 8.714 | **<0.0001** |
| | OA:Endo – OA:Blood | 10.03 | 8.003 | 12.06 | **<0.0001** |
| **$TCO_2$** | Ctrl:Endo – Ctrl:Blood | 6.256 | 4.087 | 8.425 | **<0.0001** |
| | OA:Blood – Ctrl:Blood | 2.833 | 0.7963 | 4.869 | **0.0038** |
| | OA:Endo – Ctrl:Blood | 12.84 | 10.75 | 14.94 | **<0.0001** |
| | OA:Blood – Ctrl:Endo | -3.423 | -5.535 | -1.311 | **0.0007** |
| | OA:Endo – Ctrl:Endo | 6.588 | 4.419 | 8.757 | **<0.0001** |
| | OA:Endo – OA:Blood | 10.01 | 7.975 | 12.05 | **<0.0001** |

**Table S6:** $Na^+/K^+$-ATPase (NKA) and V-type $H^+$-ATPase (VHA) protein abundance in the inner ear organ of control and OA-exposed rockfish. Values are mean ± s.e.m. There were no significant differences for NKA or VHA (Two-tailed t-test, α = 0.05).

| Protein | Relative Protein Abundance | n | t-ratio | Degrees of freedom | ***p*-value** |
|---|---|---|---|---|---|
| **NKA** | Control: 1.00 ± 0.12 | 7 | 0.11728 | 13 | 0.9104 |
| | OA: 0.98 ± 0.15 | 8 | | | |
| **VHA** | Control: 1.00 ± 0.08 | 8 | 1.4182 | 13 | 0.1695 |
| | OA: 0.78 ± 0.13 | 7 | | | |

**Table S7:** Relative comparison of $Na^+/K^+$-ATPase (NKA) and V-type $H^+$-ATPase (VHA) signal intensity and subcellular localization in rockfish inner ear epithelial cells. There were no apparent differences between control and ocean acidification-exposed rockfish.

| Cell Type | NKA Signal | VHA Signal |
|---|---|---|
| Type-I Ionocyte | +++<br>Basolateral membrane infoldings | +<br>Cytoplasm |
| Type-II Ionocyte | - | +<br>Cytoplasm |
| Sensory Hair Cell | ++<br>Basolateral membrane | +++<br>Cytoplasm (concentrated in the basal portion) |
| Supporting Cell | - | +<br>Cytoplasm |
| Granular Cell | +<br>Lateral membrane | +<br>Cytoplasm |
| Squamous Cell | ++<br>Ribbon-like lateral membrane | +<br>Cytoplasm |
